# Cytosine Methylation is a marker of Viral Gene Transfer across the eukaryotes

**DOI:** 10.1101/2025.04.15.648976

**Authors:** Luke A. Sarre, Giselle Azucena Gastellou Peralta, Pedro Romero Charria, Vladimir Ovchinnikov, Alex de Mendoza

**Affiliations:** School of Biological and Behavioural Sciences, Queen Mary University of London, London, United Kingdom; Centre for Epigenetics, Queen Mary University of London, London, United Kingdom; Wellcome Sanger Institute, Hinxton, United Kingdom

## Abstract

Cytosine DNA methylation patterns vary widely across eukaryotes, with its ancestral roles being understood to have included both transposable element silencing and host gene regulation. To further explore these claims, in this study, we reevaluate the evolutionary origins of DNA methyltransferases and characterise the roles of cytosine methylation on underexplored lineages, including the amoebozoan *Acanthamoeba castellanii*, the glaucophyte *Cyanophora paradoxa*, and the heterolobosean *Naegleria gruberi*. Our analysis of DNA methyltransferase evolution reveals a rich ancestral eukaryotic repertoire, with several eukaryotic lineages likely subsequently acquiring enzymes through lateral gene transfer (LGT). In the three species examined, DNA methylation is enriched on young transposable elements and silenced genes, suggesting an ancestral repressive function, without the transcription-linked gene body methylation of plants and animals. Notably, the closest homologues of many of the silenced, methylated genes in diverse eukaryotes belong to viruses, including giant viruses. Given the widespread occurrence of this pattern across diverse eukaryotic groups, we propose that cytosine methylation was a silencing mechanism originally acquired from bacterial donors which was used to mitigate the expression of both transposable and viral elements, and that this function may persist in creating a permissive atmosphere for LGT in diverse eukaryotic lineages. These findings further highlight the importance of epigenetic information to annotate eukaryotic genomes, as it helps delimit potentially adaptive LGTs from silenced parasitic elements.

## Introduction

5-methylcytosine (5mC) is a widespread base modification in eukaryotes, which has evolved divergent patterns and functions across different lineages(Schmitz et al. 2019; de Mendoza et al. 2020; Sarkies 2022). In plants and animals, 5mC typically accumulates along gene bodies of constitutively expressed genes, with minimal methylation in promoter regions—a pattern referred to as Gene Body Methylation (GBM)(Feng et al. 2010; Zemach et al. 2010; Bewick and Schmitz 2017). In contrast, fungi and many algal groups predominantly accumulate 5mC in transposable elements (TEs) and transcriptionally silent DNA(Huff and Zilberman 2014; Bewick et al. 2019; Hoguin et al. 2023). Interestingly, plants and certain invertebrates possess both GBM and TE methylation(de Mendoza et al. 2019; Lewis et al. 2020), leading to the hypothesis that both functions may have been present in the last common ancestor of eukaryotes(Zemach and Zilberman 2010; Yi 2024).

Supporting this hypothesis, DNA methyltransferases (DNMTs), which catalyse 5mC, are conserved across plants and animals(Law and Jacobsen 2010). DNMT1 orthologues, known as MET1 in plants, mediate maintenance methylation at CpG dinucleotides, while DNMT3 enzymes catalyse *de novo* methylation(Lyko 2018). Plants also encode additional DNMT1 paralogues in the Chromomethylases (CMT), which methylate CHG trinucleotides (H = A, T, C); and DNMT3 paralogues in the Domains Rearranged Methyltransferases (DRM) that targets the CHH context, with both CHG and CHH methylation contexts predominantly associated to silencing(Bewick et al. 2017). In contrast, fungi and algal groups lacking GBM also typically lack DNMT3(Huff and Zilberman 2014; Bewick et al. 2019), highlighting a potential connection between the presence of DNMT3 and GBM. It is however noteworthy that GBM in plants does not directly depend on DNMT3 orthologues, as these are restricted to CHH methylation while GBM is limited to the CG dinucleotides, whereas animal DNMT3 enzymes are the main drivers of GBM through the targeting of the PWWP domain to H3K36me3 marked gene bodies.

Recent work identified that the unicellular ichthyosporean *Amoebidium* encodes both DNMT1 and DNMT3, and exhibits a GBM-like methylation pattern(Sarre et al. 2024). This raises the intriguing possibility that unicellular eukaryotes with both DNMT1 and DNMT3 could reflect the origins of GBM. However, this hypothesis remains largely unexplored. Complicating matters, the distribution of DNMTs across eukaryotes is highly dynamic, with multiple instances of DNMT loss and diversification(Ponger and Li 2005; Huff and Zilberman 2014; de Mendoza et al. 2018; Hoguin et al. 2023). Moreover, the evolutionary origins of eukaryotic DNMTs remain unclear. Unlike other core chromatin components, DNMTs are not monophyletic and lack archaeal ancestry. Instead, they appear to have arisen from ancient lateral gene transfer (LGT) events from bacteria to early eukaryotes(Jurkowski and Jeltsch 2011).

While the role of 5mC in silencing transposable elements is well-established, 5mC also plays a role in silencing viral endogenisation events(Stuhlmann et al. 1981; Blais et al. 2024; Sarre et al. 2024). This includes the endogenisation of giant viruses, whose insertions can be orders of magnitude larger than typical retrotransposons or endogenous retroviruses, encoding hundreds of genes and making them potentially harder to co-exist with native eukaryotic DNA(Moniruzzaman et al. 2020). So far, reports of giant virus endogenization marked by 5mC are limited to the moss *Physcomitrium patens*, the ichthyosporean *Amoebidium* and the amoebozoan *Acanthamoeba castellanii(Maumus et al. 2014; Lang et al. 2018; Blais et al. 2024; Sarre et al. 2024)*. Given that virus-to-eukaryote LGT has occurred repeatedly throughout eukaryotic evolution(Irwin et al. 2022), a comprehensive evaluation of the relationship between 5mC and viral gene integration across diverse eukaryotic groups remains missing. Addressing this is critical, as 5mC may have facilitated the controlled integration and domestication of viral DNA into host genomes.

To address these knowledge gaps, we present the most complete phylogenetic analysis of DNMTs to date, reconstructing the ancestral DNMT repertoire of the last eukaryotic common ancestor. Our findings reveal recurrent cases of DNMT acquisition via LGT. Additionally, we characterise the 5mC methylomes of three understudied species, demonstrating that the presence of DNMT1 and DNMT3 does not correlate with GBM. Finally, we show that in species where 5mC plays a silencing role, viral LGT genes—often of giant virus origin—are frequently enriched among methylated genes. This suggests that 5mC-mediated silencing of viral DNA may have been an ancestral function in early eukaryotes, facilitating the stepwise integration of viral sequences into eukaryotic genomes.

## Results

### Eukaryotic DNMT repertoires are shaped by a mix of ancestral genes and lineage-specific orphan DNMTs

To reconstruct the evolutionary history of cytosine DNA methyltransferases (DNMTs) in eukaryotes, we compiled a large dataset encompassing all eukaryotic supergroups, along with bacterial out-groups from previous studies(de Mendoza et al. 2018; Richter et al. 2022). It has been proposed that eukaryotic DNMTs originated from multiple bacterial donors(Jurkowski and Jeltsch 2011), with little support for a single ancestral event followed by eukaryote-specific duplications. To reassess this hypothesis, we included DNMTs from giant viruses, such as those from the *Nucleocytoviricota* and Mirusviruses, since some core eukaryotic genes have been suggested to originate from, or be co-opted by, these viral lineages(Bernabeu et al. 2024; Karki et al. 2024). We also examined Asgardarchaeota archaeal genomes, as several eukaryotic-specific proteins have been traced to this group(Eme et al. 2017).

Among eukaryotic DNMTs, DNMT1 and DNMT5 displayed complex ancestral domain architectures (**Figure 1c**), indicating early functional interactions with eukaryotic chromatin. For example, DNMT1 enzymes are characterised by the presence of Bromo Adjacent Homology (BAH) and Replication Foci Domains (RFD), both essential for maintaining 5mC during DNA replication(Lyko 2018). Even lineage-specific paralogues, such as plant chromomethylases (CMT) and fungal DIM2, retain the BAH domain (**Figure 1c**). However, domain reconfiguration occurred throughout evolution, such as the addition of a zinc finger CXXC domain in holozoans or a Chromo domain in plant CMTs. Despite these changes, DNMT1 orthologues are remarkably conserved in both length and domain structure across all eukaryotic supergroups (**Figure 1d**). Similarly, DNMT5 orthologues in euglenozoans, algae, and fungi possess an additional C-terminal SNF2 ATPase domain, though this domain has been lost in some lineages, such as dinoflagellates and other algae.

**Figure 1.**
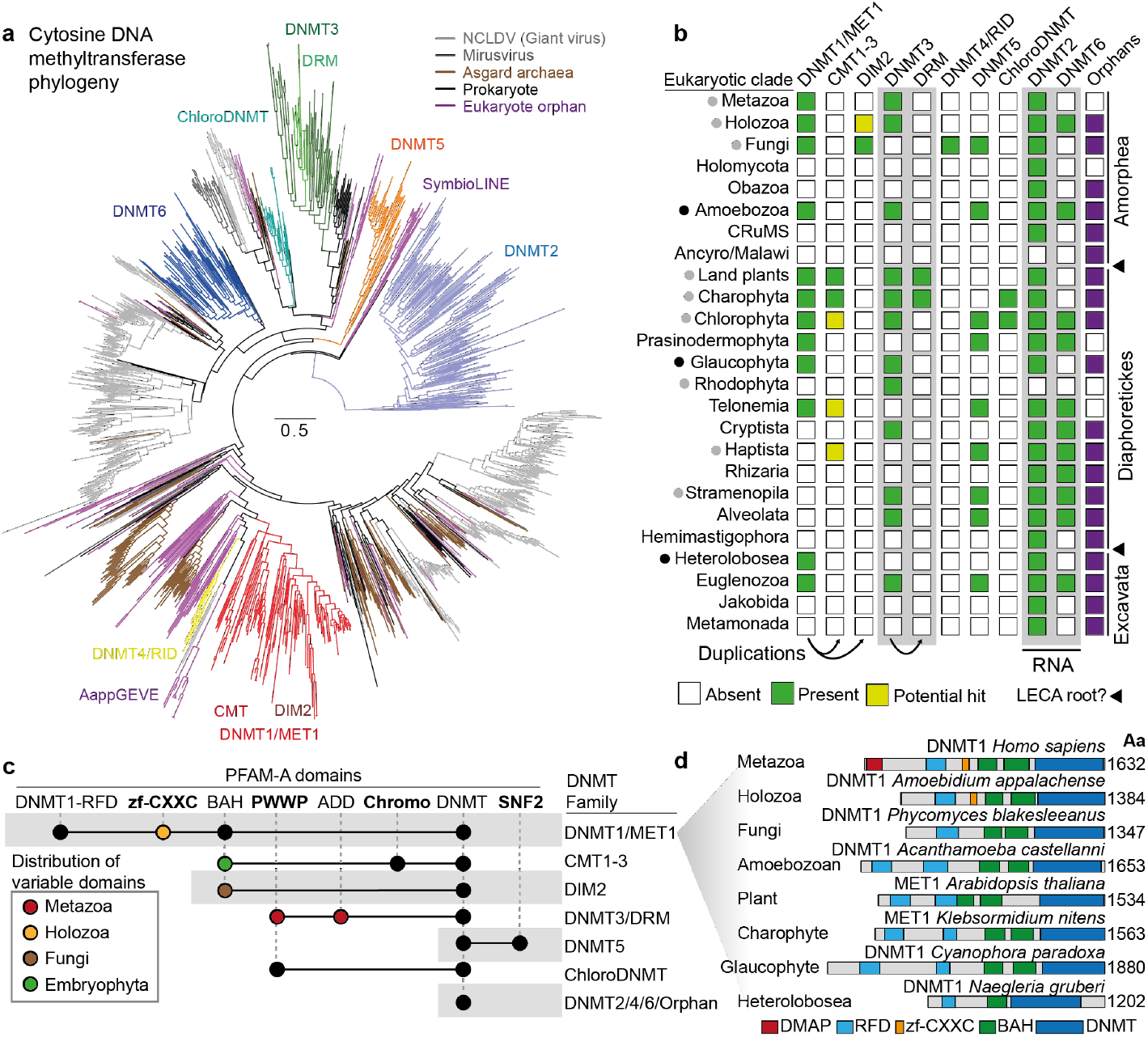
Evolutionary dynamics of DNMTs in the eukaryotes. **a**) Maximum likelihood phylogenetic tree of DNMTs, including eukaryotes, asgard archaea, viruses and bacteria. Each major eukaryotic DNMT family is highlighted with a colour. Eukaryote orphan are eukaryotic sequences that do not cluster with any previously defined family. Grey branches depict viral sequences, brown branches depict Asgardarchaeota sequences, and black sequences depict bacteria. **b**) Distribution of eukaryotic DNMTs across the eukaryotes, based on phylogenetic profiling of panel b. Green cells depict presence, whereas yellow cells represent potential presence. Potential presence implies distinct domain architectures from “archetype” and low nodal branch supports. Black triangles highlight the two competing hypotheses regarding the root of extant eukaryotes. Raw data with per species DNMT repertoires can be found in Supplementary Table 1. **c**) Protein domain architecture of the main eukaryotic DNMT families, as defined by PFAM domains. Each dot implies presence of the domain, but coloured dots indicate that the domain is restricted to an eukaryotic lineage. **d**) Domain architectures of selected DNMT1 orthologues across major eukaryotic supergroups.

In contrast, DNMT3 orthologues show little domain conservation across eukaryotes. Animal DNMT3s contain PWWP and ADD domains, land plants possess a Domain of Unknown Function(Yaari et al. 2019), and protist and algal DNMT3s lack identifiable companion domains (**Figure 1c**). DNMT2 and DNMT6, which are associated with RNA modifications(Goll et al. 2006; Raddatz et al. 2013; Cuypers et al. 2020), consistently lack any companion domains, suggesting that DNMTs involved in DNA and chromatin regulation tend to acquire multifunctional domains, while those involved in RNA methylation do not.

Beyond the eukaryotic ancestral DNMT groups, we identified several lineages with restricted distributions (**Figure 1b**). For example, DNMT4/RID appears to be confined to fungi, while a distinct lineage of DNMTs, which we term ChloroDNMTs, was found in chlorophytes and charophytes (**Figure 1b**). This group often contains a PWWP domain, likely acquired independently of animal DNMT3s. Additionally, we observed several eukaryotic DNMTs that cluster into more lineage-specific clades which we name “orphans”, some of which had been previously identified as components of dinoflagellate LINE transposable elements (SymbioLINEs)(de Mendoza et al. 2018) or as part of giant virus endogenous elements in *Amoebidium* (Figure 1a)(Sarre et al. 2024). While some of these orphan DNMTs may represent contaminants from non-axenic cultures or poorly assembled genomes, we identified high-quality genomes that suggest these DNMTs were integrated through viral sources. For instance, the amoeba *Acanthamoeba castellanii* (Neff strain) encodes a DNMT closely related to the DNMT of the giant virus *Pithovirus*. The widespread distribution of these orphan DNMTs across the eukaryotic dataset suggests that they are prone to being incorporated into eukaryotic genomes via lateral gene transfer (LGT).

Overall, our comprehensive dataset reveals that the LECA possessed a complex DNMT repertoire, including DNMT1, DNMT3, and DNMT5 as potential DNA methyltransferases, and DNMT2 and DNMT6 as RNA methyltransferases (Figure 1b).

However, the distribution of these genes across modern eukaryotes is highly uneven, with DNMT2 being the least likely to undergo secondary loss, indicating its essential role in tRNA methylation. Ancestral and lineage-specific DNMTs are found in various combinations in extant species, underscoring the dynamic evolution of DNA methylation machinery across eukaryotic lineages.

### DNMT1/DNMT3-maintained methylomes of *Acanthamoeba* and *Cyanophora* lack gene body methylation and are enriched in repressive 5mC

In plants and animals, DNMT1 and DNMT3 methyltransferases are responsible for both gene body methylation and the silencing of transposable elements. These roles have been proposed as ancestral to eukaryotes. To test this further, we expanded DNA methylation data to key eukaryotic lineages that encode both DNMT1 and DNMT3, focusing on species positioned at critical points in the evolutionary tree. Specifically, we selected *Acanthamoeba castellanii (Clarke et al. 2013)*, a free-living amoebozoan, which serves a sister group to the Opisthokonta (comprising animals and fungi), and the glaucophyte *Cyanophora paradoxa (Price et al. 2012)*, the sister group to the Viridiplantae (comprising land plants and green algae) known for their chloroplasts with ancestral cyanobacteria features.

Using Enzymatic Methyl-sequencing (EM-seq)(Vaisvila et al. 2021), we found that in *A. castellanii* and *C. paradoxa*, methylation predominantly occurs at CpG dinucleotides (1.3% and 32%, respectively, **Figure 2a, Supplementary Figure 1a**), while non-CpG dinucleotides are below the non-conversion rate (0.2-0.3%), suggesting this signal is mostly false positives. In *C. paradoxa*, typical plant methylation sequence contexts (CHG and CHH) showed methylation levels near the non-conversion rate (CHG 0.73%, CHH 0.53%), suggesting that these contexts probably evolved later with the appearance of CMT and DRM methyltransferases in streptophytes.

**Figure 2.**
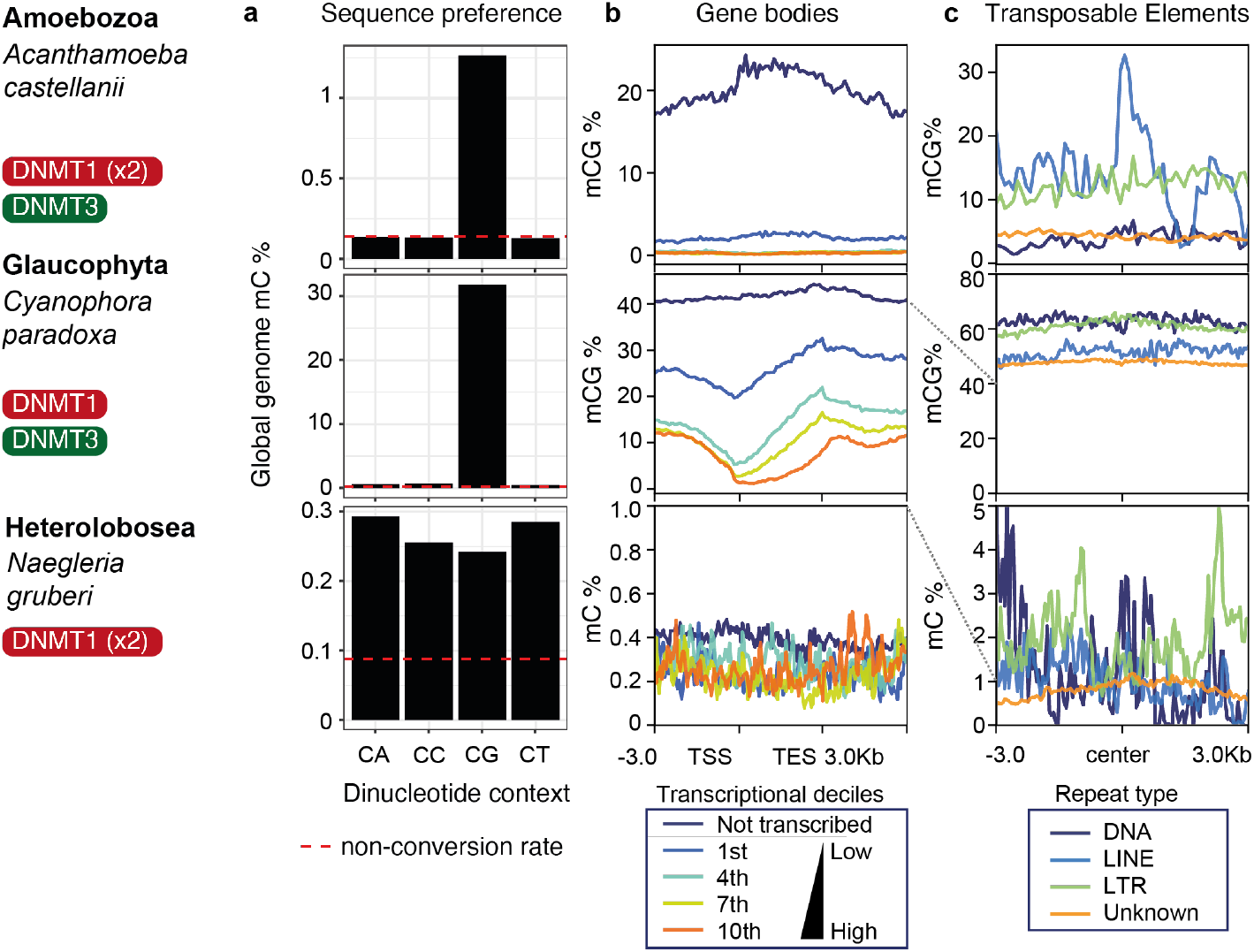
Cytosine methylome characteristics of three divergent eukaryotes. **a**) Global methylation levels at the four dinucleotide contexts as quantified with EM-seq in *A. castellanii, C. paradoxa* and *N. gruberi*. Dashed red line indicates the cytosine methylation non-conversion rate (lambda genome control) in each EM-seq experiment. b) Average methylation levels on genes separated by their transcriptional status, coloured by decile legend below. Not transcribed genes (TPM < 1). c) Average methylation levels on the three major classes of transposable elements (>500 bp) defined by RepeatModeler2, as well as unclassified repeats (Unknown), coloured by legend below.

In *A. castellanii*, methylation was enriched in repetitive and intergenic regions, while introns and exons, which make up the majority of the genome (∼80%), exhibited low methylation (**Supplementary Figure 2a**). LINE and LTR retrotransposons were preferentially methylated, particularly younger copies (estimated using Kimura distances), except for a divergent group of LTRs (**Figure 2c, Supplementary Figure 2b**,**c**). DNA transposons showed no similar enrichment. Few genes (381, 2.4%) were highly methylated (mCG ≥20%), and these genes were mostly transcriptionally silent (**Figure 2b**), with methylation extending across the gene body and promoter - a pattern unlike plants and animals, where GBM correlates with transcriptional activity and promoters remain unmethylated. Methylated genes in *A. castellanii* were enriched in a whole range of functions, including kinase activity and DNA replication (**Supplementary Figure 1b**), yet these are likely reflecting the composition of giant endogenous viruses(Blais et al. 2024).

In *C. paradoxa*, we observed similar methylation patterns. Repetitive sequences were preferentially methylated, with both retrotransposons and DNA transposons showing high methylation levels (**Figure 2c**), however repeat insertion age (estimated with Kimura distance) did not show a clear correlation with 5mC, which suggests that most repeats are effectively methylated in *C. paradoxa* (**Supplementary Figure 2d**). Moreover 1231 genes (10%) showed elevated methylation (mCG >50%, Figure 2b). Like *A. castellanii*, these methylated genes were primarily silent, indicating that methylation is associated with gene repression rather than activation. However, when separating *C. paradoxa* genes by transcriptional level, most expression deciles showed some degree of methylation accumulating towards the end of the gene body, a pattern so far unreported in other eukaryotes (**Figure 2b**).

Overall, the data from *A. castellanii* and *C. paradoxa* resemble patterns in fungi and diatoms(Bewick et al. 2019; Hoguin et al. 2023; Ruggiero et al. 2023), showing that DNMT1 and DNMT3 are not predictive of GBM, and suggesting that these DNMTs were not linked to GBM in the eukaryotic ancestor. Instead, repressive 5mC, predominantly targeting repetitive and silent genes, likely represents the ancestral function of DNA methylation in eukaryotes.

### *Naegleria gruberi* displays CX methylation maintained by DNMT1 orthologues

The methylation data from amoebozoans and glaucophytes provides critical insights into the evolution of 5mC, but to fully understand early eukaryotic methylation patterns, it is essential to include discobans. Discobans occupy a key evolutionary position, either as a sister group to Diaphoretickes or to the rest of eukaryotes(Derelle et al. 2015; Al Jewari and Baldauf 2023; Williamson et al. 2024), making their methylation patterns valuable for identifying ancestral features. Among discobans and other members of the Excavata supergroup, only a few species have retained DNMTs, with euglenozoans and heteroloboseans being one of the few encoding DNMT1 or DNMT5 orthologues. We focused on the heterolobosean *Naegleria gruberi*, a free-living amoeboid which contains a well-conserved DNMT1 enzyme and can be cultured axenically(Fritz-Laylin et al. 2010).

Using EM-seq on *N. gruberi*, we found low genome-wide methylation levels (0.28%), though these were still above the non-conversion rate (∼0.1%). Unlike *A. castellanii* and *C. paradoxa, N. gruberi* showed no preference for CpG dinucleotides—all four dinucleotide contexts were methylated at similar levels (**Figure 2a**). The symmetry of CpG methylation was modest (r = 0.25), and few individual cytosine sites were fully methylated, suggesting variability in methylation from cell to cell (**Supplementary Figure 1a**). This makes the *N*.

*gruberi* methylome particularly interesting, as it indicates that despite possessing only a single DNMT1 enzyme, the species can methylate non-CpG sites and likely exhibits *de novo* methylation activity, since the intermediate levels of cytosine methylation suggest a dynamic process of 5mC gain and loss. If this dual functionality and substrate plasticity of *N. gruberi* DNMT1 is ancestral or a lineage-specific variation is unclear.

Though 5mC was sparse in the *N. gruberi* genome, it accumulated disproportionately in repetitive regions (**Supplementary Figure 2a)**. A small fraction of these repeats could be classified as known transposable elements, with both DNA transposons and retrotransposons showing higher methylation than the genomic background (**Figure 2c**). Consistent with *A. castellanii*, younger transposable element insertions had higher methylation levels than older ones (**Supplementary Figure 2e**).

In terms of gene methylation, only 115 genes (0.6% of annotated genes) showed significant methylation (mC ≥10%), most of which were transcriptionally silent, yet with some exceptions (**Supplementary Figure 3**). Methylated genes were enriched in functions associated with cytoskeleton and translation blocking among others, yet a particular pathway was not evident (**Supplementary Figure 1b**). This finding suggests that, similar to other eukaryotes, 5mC in *N. gruberi* primarily acts as a repressive mark, reinforcing the idea that ancestral eukaryotic methylation was largely restricted to repetitive elements and silenced genes.

### Cytidine analogues partially deplete 5mC in *A. castellanii*

To directly assess the repressive function of 5mC in *A. castellanii* and *N. gruberi*, we treated cells with cytidine analogues known to incorporate into DNA and inhibit DNMTs, leading to passive 5mC dilution. We applied 5-Azacytidine, which is incorporated into both RNA and DNA in mammals, and its derivatives, Decitabine and Zebularine which are DNA-specific (Gowher and Jeltsch 2004). DMSO was used as a control, since it is required to solubilize the inhibitors. No growth defects were observed at any concentration. We selected concentrations that stayed below 1% DMSO, as higher levels affect *A. castellanii* growth (Siddiqui et al. 2016), yet were effective at demethylating based on concentrations previously used in the protist *Amoebidium (Sarre et al. 2024)*.

After 3 days of treatment, we used EM-seq to profile methylation levels in both species. In *A. castellanii*, Decitabine and 5-Azacytidine markedly reduced global mCG levels by 29.5% and 23.4%, respectively, while Zebularine had no effect (**Supplementary Figure 4a**,**b**). In contrast, no inhibitors reduced methylation in *N. gruberi* (**Supplementary Figure 4e**), highlighting that cytidine analogues vary in efficacy across species, with some exhibiting the expected demethylation and others showing no response, which recapitulates previous observations in animals.

RNA-seq analysis revealed a broad transcriptional response in *N. gruberi* to cytidine analogues, with hundreds of genes differentially expressed across treatments, although only a fraction of methylated genes were affected, and these changed expression at lower fold changes than many unmethylated genes (**Supplementary Figure 4f,g**). Interestingly, both 5-Azacytidine and Decitabine led to comparable gene expression changes (**Supplementary Figure 4f**), predominantly affecting unmethylated genes associated with RNA export and protein polymerization (**Supplementary Figure 5**), suggesting that the cytidine analogues can exert broad transcriptional effects independent of DNA methylation.

In *A. castellanii*, three methylated genes were consistently upregulated upon demethylation treatment, ranking among the top differentially expressed genes sorted by transcriptional fold change (**Supplementary Figure 4c**). However, many unmethylated genes were differentially regulated with 5-Azacytidine or Decitabine, showing both up-regulation and downregulation upon treatment (**Supplementary Figure 4d**), suggesting widespread effects on the transcriptome beyond methylated genes, affecting various processes including cytoskeleton organisation to metabolic functions (**Supplementary Figure 6**). In fact, the vast majority of the 381 methylated genes remained transcriptionally inactive. Among the repetitive elements that were upregulated, none were initially methylated. This indicates that cytidine analogues only partially inhibit methylation in *A. castellanii* and produce scarce transcriptional reactivation in methylated regions, while having substantial off-target effects. Since *A. castellanii* is known to be polyploid (Matthey-Doret et al. 2022), perhaps the methylation loss is not effective enough to robustly reactivate transcription across all chromosome copies, or maybe there are other regulatory mechanisms (e.g. heterochromatin-compacted DNA) preventing transcriptional reactivation of methylated genes and TEs.

Among the methylated genes that were consistently upregulated in *A. castellanii*, all were part of giant virus endogenised regions(Blais et al. 2024). This suggests that these few genes from viral origin might be either especially sensitive to DNA methylation loss, or might be responsive to a non-specific stress response caused by cytidine analogues.

### Genes from viral origins are frequently methylated in eukaryotic genomes

In most eukaryotic lineages with the notable exceptions of plants and animals, methylated genes are transcriptionally silent, belonging to transposable elements open reading frames, or part of hypermethylated giant virus endogenised elements (GEVEs). If 5mC is used as a mechanism to recognise and silence non-self DNA across eukaryotes, we wondered if the methylated fraction of genes would reveal the parts of the genome that have potential non-eukaryotic origins across several lineages.

To test this, we used the data for *A. castellanii, C. paradoxa* and *N. gruberi* and gathered whole genome methylation data from a subset of eukaryotes for which 5mC is associated with silencing (de Mendoza et al. 2018; Griess et al. 2023; Hoguin et al. 2023; Sarre et al. 2024). For each species, we performed a DIAMOND search against NCBI non-redundant database, gathering the ten best hits for each gene and assigning potential taxonomic origin. In parallel, for each gene we classified them as methylated or unmethylated. The inclusion in the methylation category was tailored to each species particular methylome patterns, focusing on genes methylated in the sequence context that is associated with silencing in a given species (e.g. CHG methylation in streptophytes, CG in *A. castellanii*, or CX in *N. gruberi*).

Crossing the information from potential taxonomic origin and methylation status, we observed that the methylated fraction tends to be enriched in genes with potential recent viral ancestry (**Figure 3a, Supplementary Figure 7**). While *A. castellanii*, two *Amoebidium* species (*appalachense* and *parasiticum*), *C. paradoxa, Klebsormidium nitens*, and *Physcomitrium patens* showed strong methylation enrichments for the viral fraction of the genome, other species were less evident, for instance, we did not see an enrichment for viral genes in *N. gruberi* or its sister species *N. fowleri* (Liechti et al. 2019), and for the diatom *Phaeodactylum tricornutum*, where the only 2 genes in the genome with potential viral ancestry were methylated, but were insufficient to render a significant enrichment (**Figure 3a, Supplementary Figure 7**). We further searched the taxonomic origin of methylated genes in several fungal species described to have islands of methylated genes (Bewick et al. 2019), and we did not find any support for those clusters of methylated genes having viral origins. In contrast, in most species methylated genes with eukaryotic origins were overwhelmingly genes with transposable element domains. Noteably, the fraction of genes from potential prokaryotic LGTs were less likely to be enriched in the methylated fraction (**Figure 3a**). This implies that these genes might be less prone to silencing by 5mC than genes from viral origin.

**Figure 3.**
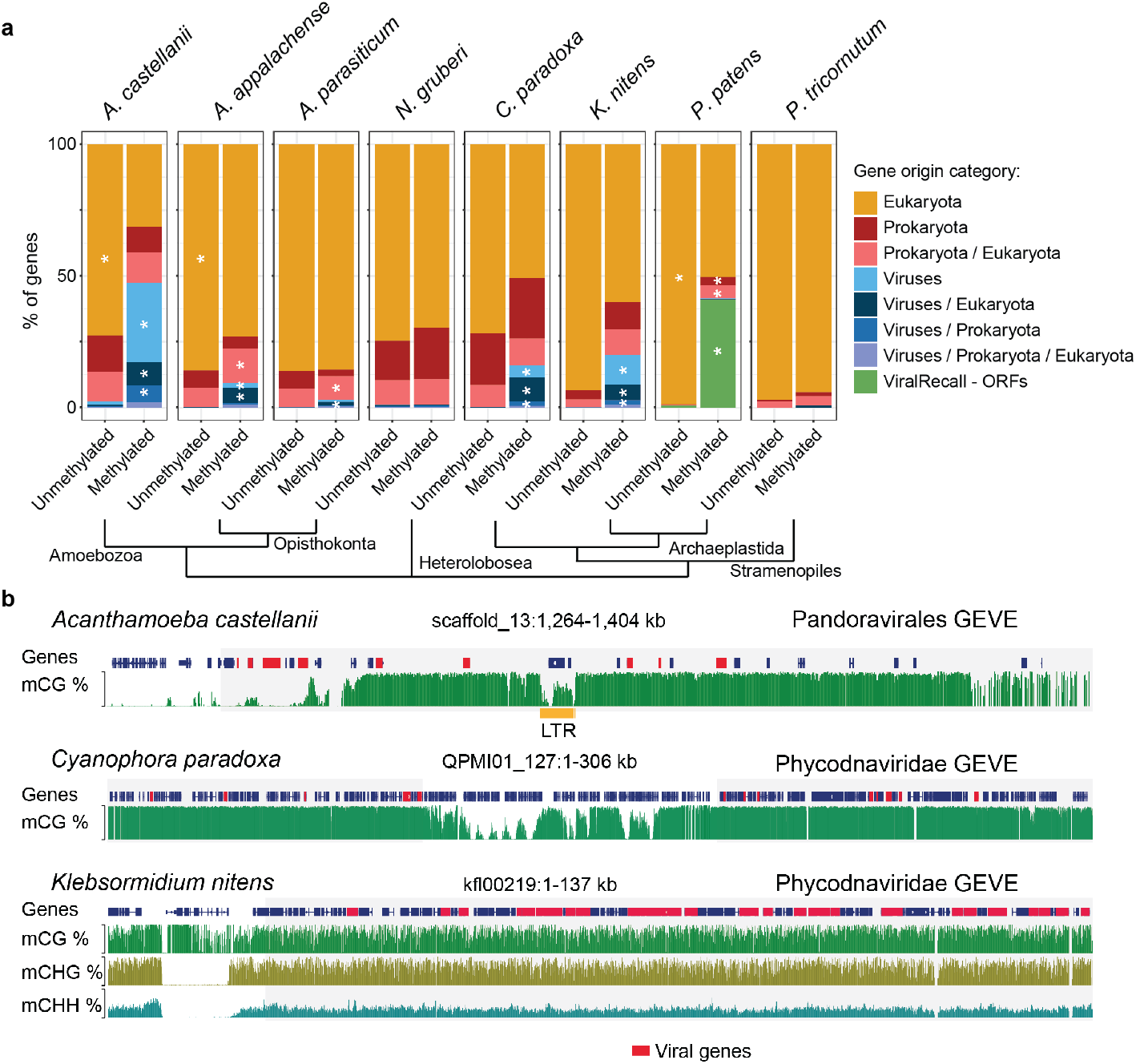
Cytosine methylation is enriched on genes from viral origin across divergent eukaryotes. **a**) The proportion of genes based on potential phylogenetic origins (best hits against NCBI nr database, see Methods) per species, split by methylation status. For *P. patens*, we ran ViralRecall as all viral genes have been masked from the Phypa V5 genome version. For *A. castellanii* methylated genes are defined as mCG ≥ 20%, in *Amoebidium appalachense* / *parasiticum* as non-mCGC methylation ≥ 70%, for *N. gruberi* mC ≥ 5%, for *C. paradoxa* mCG ≥ 50 %, for *K. nitens* and *P. patens* mCHG ≥ 10 %, for *P. tricornutum* ≥ 1 %. White asterisks represent statistical enrichment (two-sided Fisher exact test < 0.01). **b**) Genome browser examples of Giant Viral Endogenous Elements (GEVEs) identified in *A. castellanii, C. paradoxa* and *K. nitens* genomes. Potential donor NCLDV lineages depicted on the top right as per best hits against NCBI nr database, genes with closest viral gene origins depicted in red. Yellow box indicates LTR insertion.

While the genes of recent viral ancestry mapped to a wide variety of viruses across the different species, the overwhelming majority were assigned to giant virus (*Nucleocytoviricota*) origin. When inspecting the genomic location of these genes, they tend to cluster in genomic neighbourhoods. This approach revealed GEVEs in species for which these were not yet well described, including *C. paradoxa* or *K. nitens* (**Figure 3b**). This expands the presence of GEVEs to branches of the tree of life for which these were not yet reported, and highlights how DNA methylation information can be very useful to determine the origin of important parts of the coding potential of a given genome.

## Discussion

Through profiling the 5mC methylomes of three eukaryotes at key phylogenetic positions of the eukaryotic tree of life, we demonstrate that 5mC silencing of transposable elements and viral-derived genes is widespread across eukaryotes. Yet, despite its potentially ancestral repressive function, 5mC patterns are highly dynamic and have diversified significantly over evolutionary time. In most eukaryotes, including the three species studied here, 5mC is predominantly associated with silenced genomic regions (de Mendoza et al. 2020). However, the GBM typical of plants and animals - where gene bodies are methylated but promoters remain unmethylated, often correlating with active transcription - is relatively rare and likely evolved convergently in these multicellular lineages (**Figure 4**). An important difference between these two patterns is that in plants and animals, genes with GBM tend to be highly conserved (Sarda et al. 2012; Takuno and Gaut 2013; Guynes et al. 2024), whereas in species with 5mC playing exclusive repressive roles, methylated genes tend to be poorly conserved, even at the species level as seen across *Acanthamoeba* strains(Blais et al. 2024). Then, some unicellular lineages have large fractions of genes with methylation along the whole gene body, but this is unrelated to transcription, and presents many lineage-specific variations, such as default hypermethylation of promoters and gene bodies in dinoflagellates(de Mendoza et al. 2018; Li et al. 2024), or internucleosomal methylation in haptophytes and prasinophytes(Huff and Zilberman 2014). Despite the ancient conservation of DNMT families, predicting 5mC patterns just from DNMT repertoire remains elusive. Here we find that some DNMT3 encoding eukaryotes do not show GBM. Similarly, not all species whose methylomes are regulated by DNMT5, such as prasinophytes and haptophytes, present a distinctive internucleosomal 5mC pattern in gene bodies (Huff and Zilberman 2014). Species like the diatom *Phaeodactylum tricornutum* and the fungus *Cryptococcus neoformans* encode DNMT5 but exhibit a more restrictive methylation pattern limited to transposable elements (Huff and Zilberman 2014; Catania et al. 2020; Hoguin et al. 2023). This variation underscores the complexity and diversity of 5mC function across eukaryotes. Despite expanding our understanding, a vast portion of eukaryotic methylation remains uncharacterized, particularly among unicellular eukaryotic lineages. Advances in sequencing technology, particularly long-read methods like Oxford Nanopore and PacBio, which can natively detect 5mC, promise to rapidly expand our knowledge of methylation across new eukaryotic genomes. From our survey of DNMTs, some particularly promising lineages that deserve attention are the Euglenids, *Prasinoderma*, charophytes or early branching brown algae encoding DNMT1 orthologues(Denoeud et al. 2024). However, a major challenge remains: the frequent loss of DNMTs in various eukaryotic groups. There is ample evidence that species only encoding RNA methyltransferases (DNMT2 or DNMT6) or lacking DNMTs, have undetectable 5mC levels(Raddatz et al. 2013; Cuypers et al. 2020; Kyger et al. 2021; Drewell et al. 2023; Sarre et al. 2024), which is important when considering 5mC profiling in new lineages, especially since most 5mC techniques have some level of false positives. This evolutionary trend toward a reduced DNMT repertoire suggests redundancy among DNMTs, as no extant species retains the full complement of DNMTs we hypothesise were present in the LECA (**Figure 4**). For instance, the presence of the maintenance enzyme DNMT1 might render DNMT5 redundant, and vice versa. In stark contrast, the green lineage, particularly the streptophytes, exhibits a complex expansion of its DNMT repertoire, incorporating DRM and CMT methyltransferases, allowing for methylation in broader sequence contexts (CG, CHG, and CHH), often tied to small RNA-mediated defence mechanisms(Yaari et al. 2019). This complex DNMT repertoire is rarely lost in plants, hinting at a conserved evolutionary pressure for maintaining these mechanisms, which might have evolved from simpler patterns like those we observed in *C. paradoxa*. However, even in plants some exceptions exist, as loss of GBM has been reported in at least two angiosperms (Bewick et al. 2016; Muyle and Gaut 2019). However, total loss of 5mC is rampant across the eukaryotes, which is likely due to its potential cytotoxic effects and mutagenic potential (Pfeifer 2006; Rošić et al. 2018).

**Figure 4.**
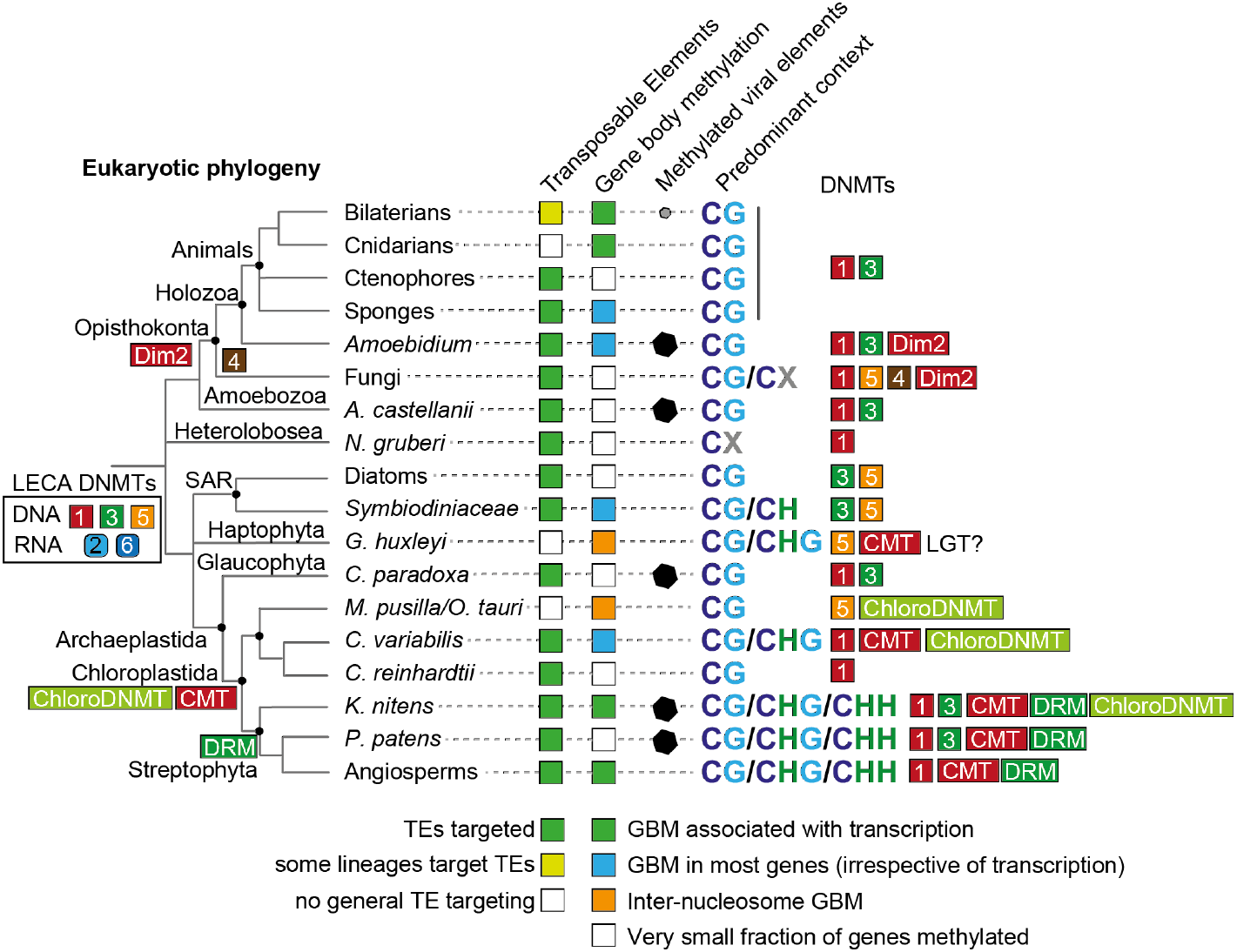
Diagram of 5mC pattern evolution across the eukaryotes. Phylogenetic tree of eukaryotes is based on current consensus branching patterns, and main DNMT families are shown as numbered coloured squares, showing gains in the branches where they appear. Only lineages for which direct whole genome 5mC data is available are shown. TE and GBM are subdivided as different cases, specified in the legend below. Methylated viral elements are shown for confirmed cases, in the case of bilaterians, it is restricted (so far) to endogenous retrovirus in vertebrates. Predominant 5mC sequence context in the lineage, while some exceptions exist (e.g. CH methylation in the vertebrate neural system(de Mendoza et al. 2021)). Main DNMTs found in each lineage, although secondary loss occurs in many instances (e.g. many animals have lost DNMT1/3).

Beyond the role of 5mC in transposable element silencing, our findings also highlight its crucial role in the epigenetic repression of viral endogenisations. Viral sequences can represent a significant proportion of eukaryotic genomes, yet are rarely expressed. Our results suggest that, at least in species where 5mC acts as a repressive mark, viral-derived genes are frequently hypermethylated, likely as a mechanism to silence these potentially detrimental sequences. This raises intriguing evolutionary questions. While LGT is predicted to be rare in eukaryotes outside of major endosymbiotic events (Ku et al. 2015), frequent reports of viral-derived sequences in eukaryotic genomes nonetheless shows its importance (Irwin et al. 2022). Epigenetic silencing through 5mC provides a mechanism by which foreign DNA, especially viral sequences, may be integrated into host genomes without imposing immediate fitness costs (Blais et al. 2024; Sarre et al. 2024). This reconciles the apparent contradiction between the scarcity of conserved LGT in eukaryotes and the frequent detection of viral sequences in their genomes. However, viral endogenisations are not necessarily evolutionarily neutral. While most viral-derived genes are transcriptionally silent and under epigenetic control, some may occasionally gain a functional role, much like endogenous retroviruses in mammals, where viral-derived genes (e.g., syncytin) or regulatory elements have been co-opted for essential biological functions(Feschotte and Gilbert 2012). These viral domestication events, though relatively rare, highlight the potential for viral endogenisations to contribute to host adaptation(Johnson 2019). However, caution must be taken in ascribing function to endogenous viral DNA, as genome-wide LGT detection methods only relying on sequence comparisons and phylogenetic profiling can often overestimate functional significance of LGT by failing to account for the silenced state of many viral-derived genes. The methylated fraction of the genome is less likely to be conserved across species or isolates(Blais et al. 2024), contributing to the non-essential pangenome.

In conclusion, our study provides further evidence that 5mC serves as a critical epigenetic gatekeeper, regulating the incorporation and silencing of foreign DNA in eukaryotic genomes. We propose this is a potentially ancestral role of 5mC in the LECA, or at least it has evolved convergently across many lineages. As we continue to map out 5mC patterns across diverse lineages, it becomes clear that epigenetic silencing is a key process allowing eukaryotes to integrate and control viral and transposable element insertions, contributing to the dynamic landscape of eukaryotic genome evolution.

## Material and Methods

### DNMT search and phylogenetic analysis

We gathered a dataset comprising a wide range of eukaryotic proteomes using EukProt database(Richter et al. 2022), complemented with some extra eukaryotic species (see Table S1). Furthermore, we also gathered all the giant virus genomes from Giant Virus DB(Aylward et al. 2021), and all *Mirusvirus* from a previous publication(Gaïa et al. 2023). All Asgardarchaeota genomes were downloaded from NCBI. We used HMMER3 to search these proteomes using the cytosine methyltransferase domain PF00145 HMM profile from PFAM, requiring a E-value < 0.0001. The subsequent hits were extracted and all the viral and archaeal hits were collapsed using CD-HIT, clustering sequences with identity above 0.8 (-c 0.8). The resulting sequences were then aligned using MAFFT with the L-INS-i parameter(Katoh and Standley 2013), and trimmed using TrimAL with gappyout mode(Capella-Gutiérrez et al. 2009). The resulting trimmed alignment was then used as input for IQ-TREE2 allowing for automatic model testing(Minh et al. 2020).

All DNMT sequences were then searched using the hmmscan function in HMMER3 to characterise the Pfam-A domain architectures, using an E-value threshold of 0.001.

### Cell culture

*A. castellanii* Neff strain was grown at 23°C in ATCC medium (712 PYG), using 100 mL ventend flasks. *N. gruberi* NEG-M was grown at 30°C in 1034 Modified PYNFH medium. DNA from untreated cultures of *A. castellanii* and *N. gruberi* were kindly provided by Meritxell Antó of Ruiz-Trillo Lab, Institut de Biología Evolutiva, Barcelona. Purified genomic DNA from *Cyanophora paradoxa* was purchased from CCAP collection.

### Treatment with cytidine analogues

DNA demethylation drugs 5-Azacytidine (ab142744 Abcam), decitabine (ab120842), and zebularine (ab141264) were dissolved in DMSO as per manufacturer’s instructions. *A. castellanii* was grown with 0 M, 10 nM, 100 nM, 1 µM, 10 µM, and 100 µM final concentration of each drug in 4ml ATCC (712 PYG) 1% DMSO media in a 6 well plate, and effects were tracked daily for 5 days, in duplicate. No treatments displayed a growth phenotype. *A. castellanii* DNA and RNA were extracted from cultures grown for 4 days in 3.5 mL ATCC (712 PYG) 1% DMSO media, with or without their respective drug in triplicate, at a final drug concentration of 100µM. *N. gruberi* DNA and RNA were extracted from cultures grown under the same conditions, with 1034 Modified PYNFH media. DNA from drug-treated cultures were extracted using the NEB Monarch Genomic DNA Purification Kit. Total RNA from these cultures were extracted using the Monarch Total RNA Miniprep Kit.

### Methylome library construction, sequencing and mapping

For each library, 100-200 ng of initial gDNA were spiked in with unmethylated lambda genome control and methylated pUC19 plasmid control. These were sheared to 300 bp using a Covaris sonicator, and then used as input for Enzymatic Methyl-Seq kit (New England Biolabs), following manufacturer instructions. The resulting libraries were sequenced at NovoGene, using Illumina NovaSeq6000 platform, aiming at coverages >30x for reference methylomes, and 5x for shallow treatment samples. The obtained FASTQ files were mapped to the reference genomes(Price et al. 2019; Matthey-Doret et al. 2022; Romero Charria et al. 2024) using BS-Seeker2(Guo et al. 2013), with BOWTIE2 as backend alignment tool. PCR duplicate reads were removed using Sambamba, and methylation calling was obtained using CGmapTools(Guo et al. 2018). The resulting CGmap files were then imported into R to be analysed using the bioconductor package bsseq. Non-conversion rates for all experiments were obtained computing all C methylation percentage on the lambda genome (∼0.3% in all experiments), while methylation rate at all the CpGs in the pUC19 plasmid were used to control for false negatives (∼98% mCG). Bigwigs of CG and CX methylation were generated using bedGraphToBigWig command from UCSC tools, and then used as input for deepTools(Ramírez et al. 2014) to produce average methylation plots and visualize in the IGV genome browser.

Repeat libraries were obtained from previous studies(Romero Charria et al. 2024), and are based on RepeatModeler2 with LTR module activated(Flynn et al. 2020), then mapped against the genome using RepeatMasker with default parameters. The Kimura distances for each transposable element insertion was obtained using the RepeatMasker alignments extracting divCpGMod values.

Publicly available Whole Genome Bisulfite libraries were downloaded from NCBI (SRX3236100, SRX22725540, SRX14311868, SRX12841637, SRR042643, SRR042657,)(Zemach et al. 2010; de Mendoza et al. 2018; Bewick et al. 2019; Griess et al. 2023; Hoguin et al. 2023; Sarre et al. 2024) and analysed using the same pipeline.

Genes were assigned a status of “methylated” or “unmethylated”, using a threshold level of cytosine methylation within gene bodies. This threshold was chosen to bifurcate bimodal distributions of gene body methylation levels where possible. No strong sequence-dependent methylation patterns were observed in *N. gruberi*, and so all cytosines were used to calculate methylation levels. In other species, methylation levels were calculated using cytosines in differing nucleotide contexts depending on the genome-wide methylation patterns. *A. castellanii, C. paradoxa*, and *P. tricornutum* almost exclusively possess methylation in the mCG context which was therefore used to calculate methylation levels, while methylation in non-CGC was previously found to mark silenced regions in *Amoebidium* species and mCHG was found to do so in *K. nitens* and *P. patens*. In these species these respective contexts were therefore used to calculate methylation levels.

### RNA-seq library construction, mapping and analysis

We used 200 ng of RNA from treated samples to build mRNA-seq libraries, first enriching for poly-A transcripts with the NEB Magnetic mRNA Isolation Kit S1550S, and then building the libraries with the NEBNext® Ultra™ II Directional RNA Library Prep Kit for Illumina® (E7760L) according to manufacturer’s instructions. Short read Illumina reads were obtained with a NovaSeq6000 at NovoGene. For *C. paradoxa*, we used publicly available RNA-seq data (SRR8306033)(Gould et al. 2019).

We then used HISAT2 to map RNA-seq reads against the reference genomes(Sirén et al. 2014), to obtain protein coding gene TPMs using Stringtie, and to perform differential expression analysis of transposable elements and protein-coding genes we obtained counts with the TElocal pipeline(Jin et al. 2015). The counts were then analysed with DEseq2 (Love et al. 2014) with default parameters, using DMSO vs treatment comparisons and selecting for genes with a p-adjusted value < 0.05 as differentially expressed. Only transposable elements above 500 bp were kept for the analysis to avoid spurious small repeat annotations. Gene Ontology enrichments were obtained with the TopGO Bioconductor package, using eggNOG-mapper annotations as input(Cantalapiedra et al. 2021).

### Proteome taxonomic-origin assignment

The proteome for each genome annotation was filtered to obtain only the longest / primary isoform per gene, and then DIAMOND (v2.1.9.163(Buchfink et al. 2021)) was used to search hits against NCBI non-redundant database (version April 2024), obtaining 10 best hits per sequence with an e-value cutoff of 0.001. The hits were then assigned taxonomic values using the diamond_add_taxonomy tool (https://github.com/pvanheus/diamond_add_taxonomy). Within-genus hits were discarded from the resulting list (e.g. *Acanthamoeba* hits against *A. castellanii*). The distribution of hits was used to obtain taxonomic categories at the highest level (Eukaryotic, Prokaryotic, Viral and combinations). For *P. patens* genome(Lang et al. 2018), the GEVEs are masked in the V5 genome annotation, therefore we used ViralRecall(Aylward and Moniruzzaman 2021) with default values to obtain ORFs of GEVE regions.

## Acknowledgements

We would like to thank Iñaki Ruiz-Trillo and Meritxell Antó for sharing the initial cultures used in this study. This work used computing resources from Queen Mary University of London’s Apocrita HPC facilities. This work was funded by the Horizon 2020 Framework Programme to AdM (European Research Council Starting Grant action number 950230), LAS and PCR were funded by a QMUL PhD fellowship, GAGP was funded by CONACyT-IPN MRes fellowship.

## Data availability

The data generated for this manuscript has been uploaded to GEO: GSE287846 (reviewer token *szgzykkanhmdjab*). Code and files to reproduce the analysis presented in this manuscript can be found on this GitHub repository: https://github.com/AlexdeMendoza/Eukaryotic_5mC_Evolution/.

## Notes

### Competing Interest Statement

The authors have declared no competing interest.

