## SupplementaryFigures for "Cytosine Methylation is a marker of Viral Gene Transfer across the eukaryotes"

Includes Supplementary Figures 1 to 7.

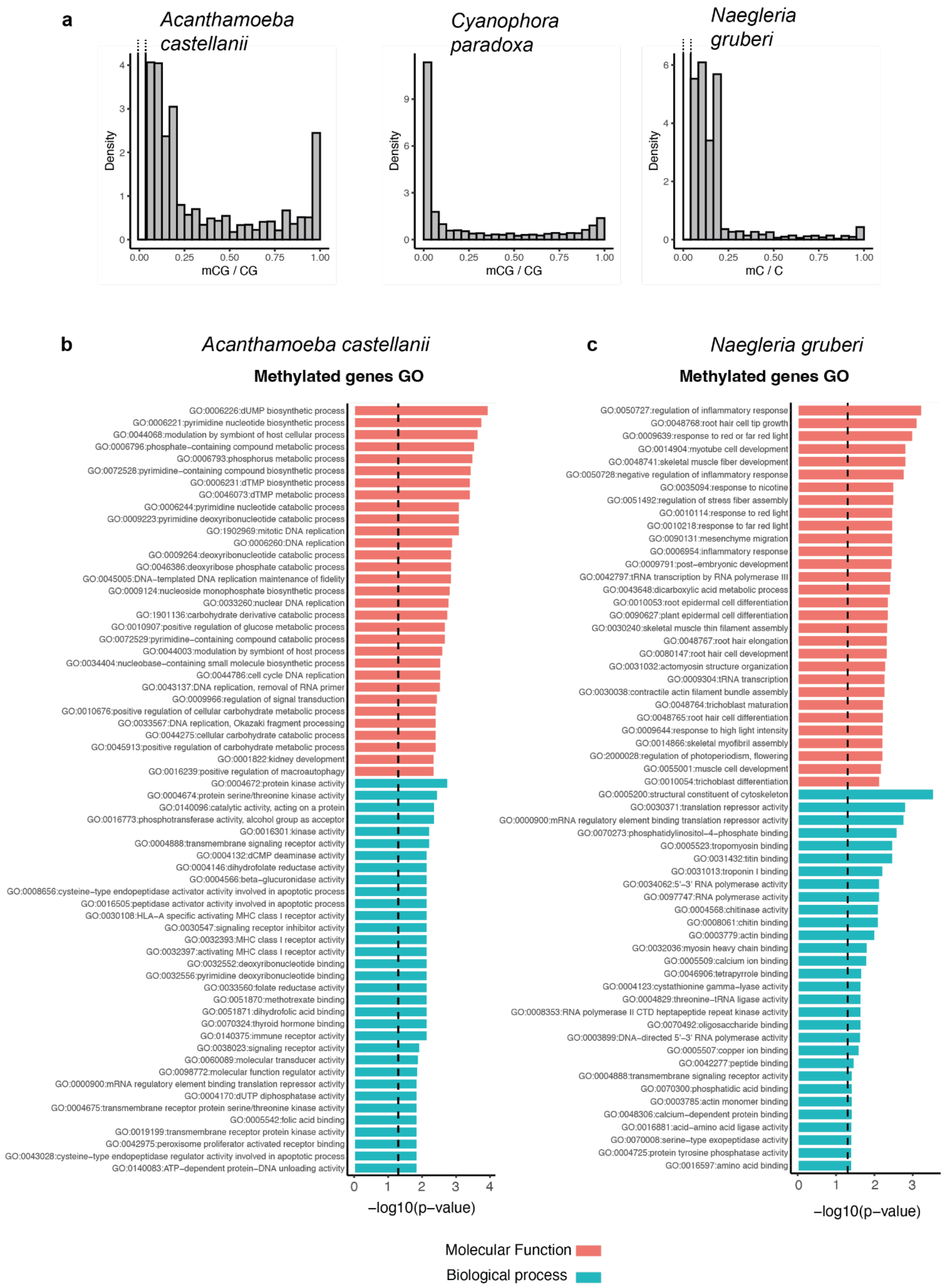


**Supplementary Figure 1. Methylation distributions on CpGs and genes**. **a**) Distribution of methylation fraction levels at all CpG sites with coverage ≥ 10x in *A. castellanii*, *C. paradoxa* and *N. gruberi* genomes. For *A. castellanii* and *N. gruberi*, sites with methylation fraction = 0 are not depicted in the same scale as these would mask the methylated site distribution. Gene Ontology enrichments sorted by significance based on Molecular Function (red) and Biological process (green) for the methylated genes in *A. castellanii* (**b**) and *N. gruberi* (**c**). Vertical dashed line indicates p-value of 0.05 according to one-sided Fisher's exact test, as calculated by TopGO.

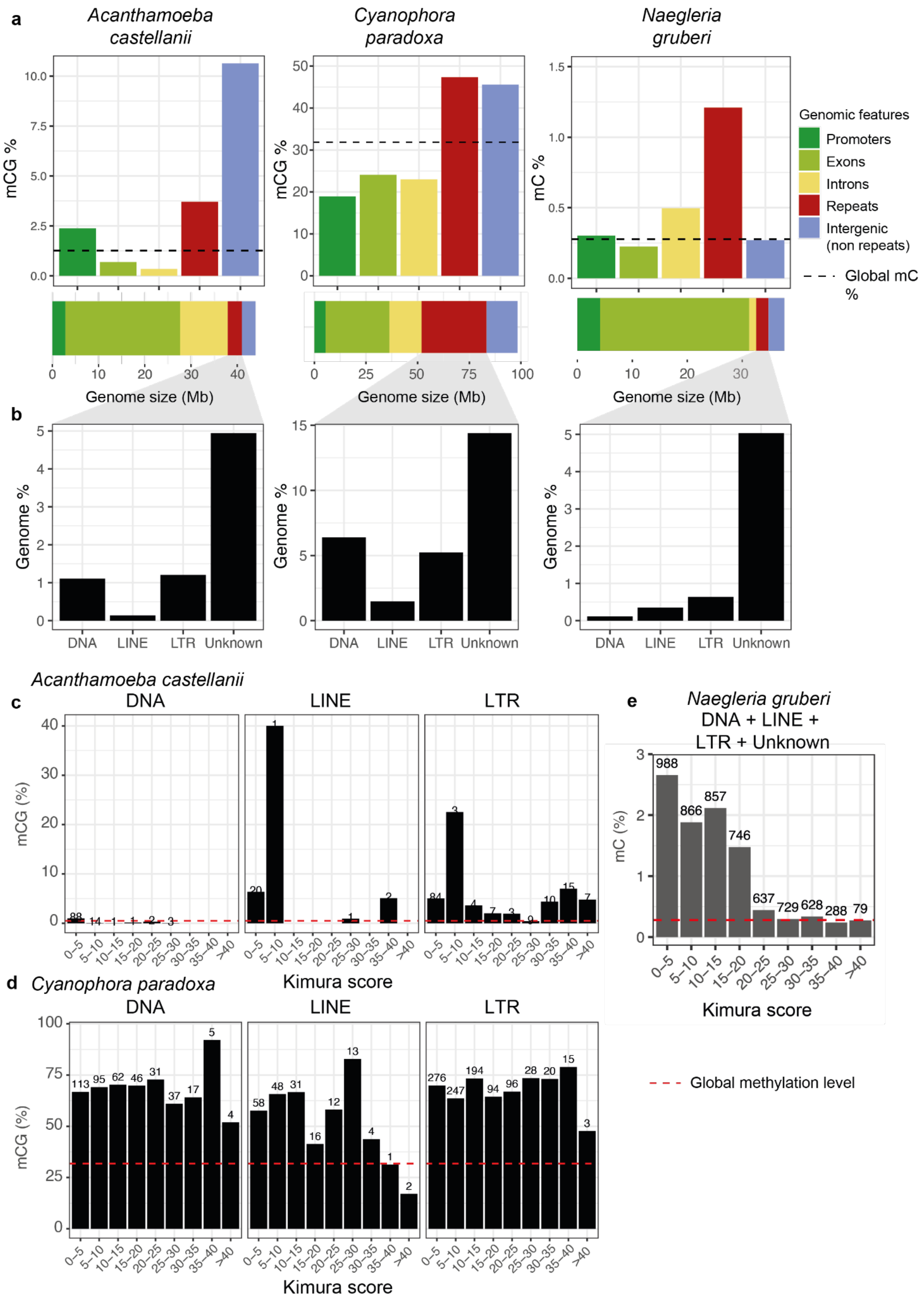


**Supplementary Figure 2. Repetitive element methylation patterns in three eukaryotes**. **a**) Global methylation levels on different features of the *A. castellanii*, *C. paradoxa* and *N. gruberi* genomes, as defined in legend. The dashed line indicates the global methylation level in the mCG, or mCX context for *N. gruberi*, displaying which features contribute disproportionately to the methylation levels. Below, the proportion of each genomic feature in each genome, shown at a scale of the total genome size. **b**) Genomic contribution of major transposable element classes and unknown repeats as defined by RepeatModeler2. **c**) Average methylation levels per each transposable element class in *A. castellanii* classified as per their Kimura divergence value, obtained from RepeatMasker. Younger elements are on the left hand side, whereas more divergent insertions are on the right hand side of each plot. Red dashed line indicate the global methylation in the genome. Numbers indicate the total amount of insertions in each Kimura range category. **d**) Same display as panel c but for *C. paradoxa*. **e**) Same display for *N. gruberi*, but combining all categories due to the very low numbers of classified transposable elements in this genome. Methylation levels as mC instead of mCG.

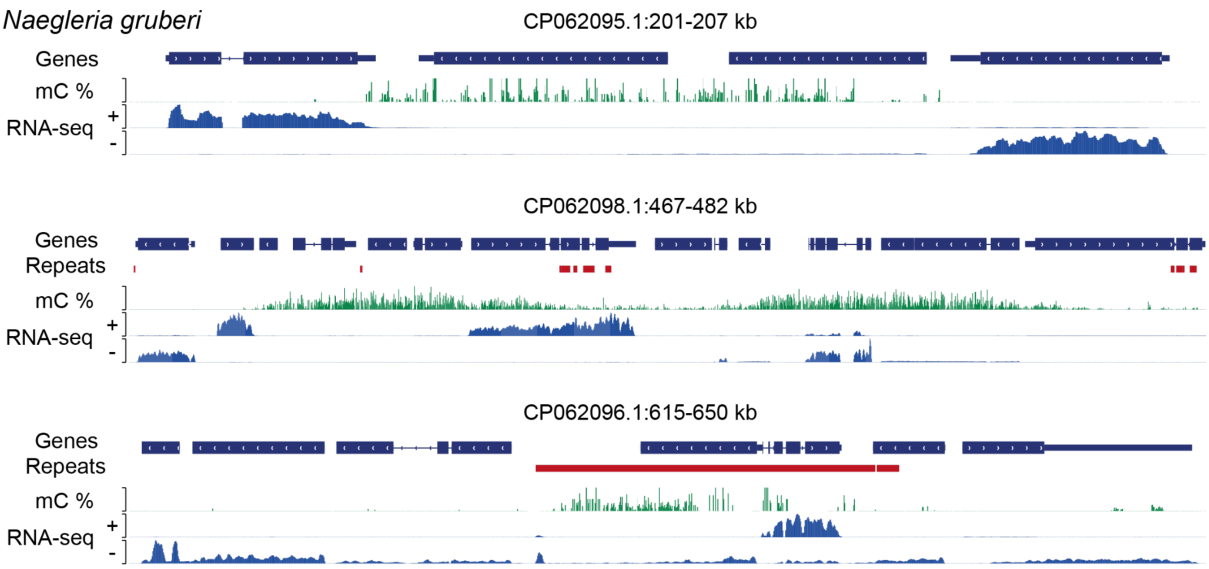


**Supplementary Figure 3. Methylated genes in the *N. gruberi* genome**. IGV genome browser representations of methylated genes / regions. Gene track includes all gene models, including UTRs and introns. In red, the RepeatMasker track of repeats, from the RepeatModeler2 library. The vast majority of repeats shown are “Unknown/Unclassified”. mC values range from 0 to 100%, whereas RNA-seq levels are split per strand. Some genes in methylated regions are actively transcribed, as shown in second and third examples.

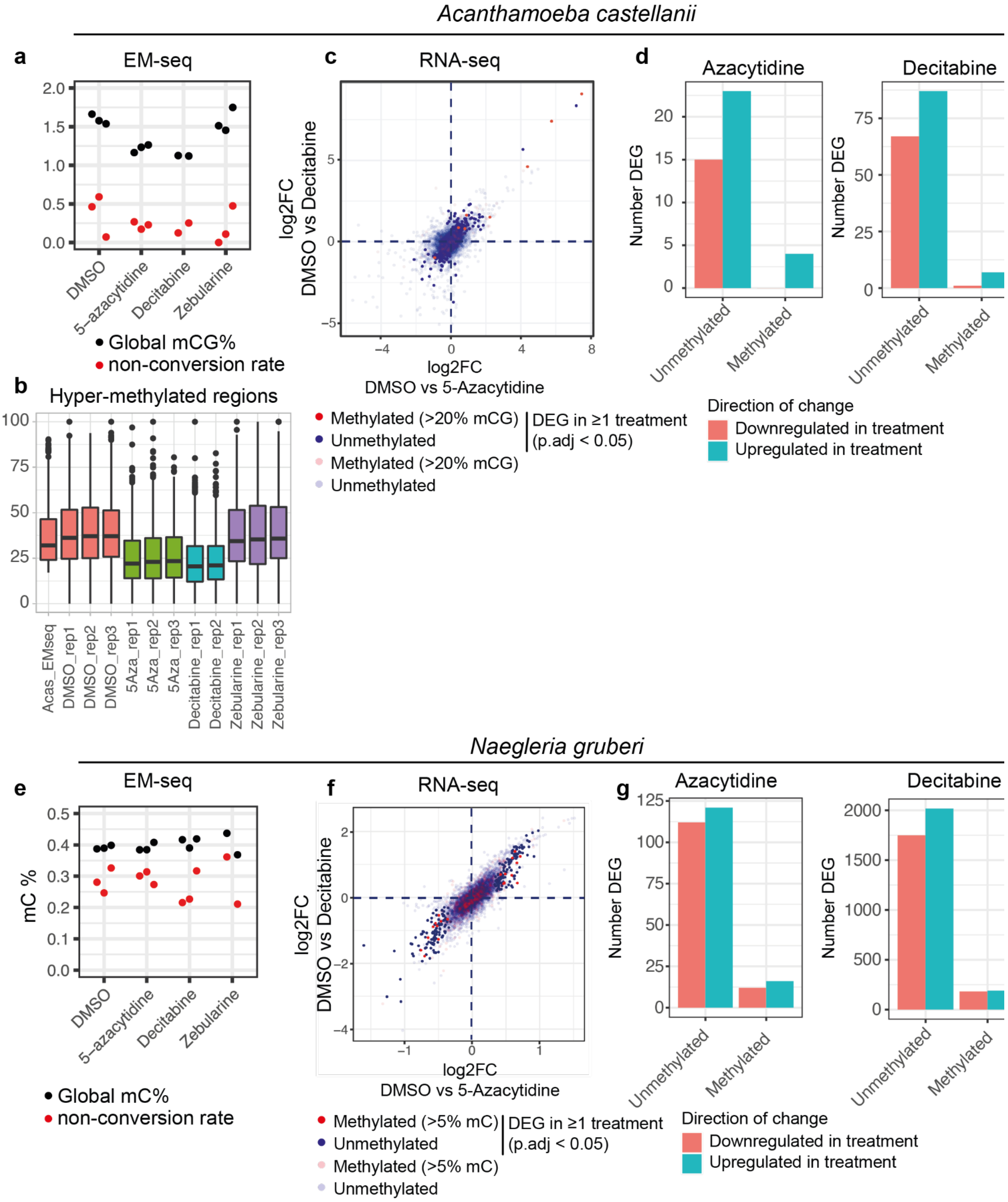


**Supplementary Figure 4. Cytidine analogues treatments on *A. castellanii* and *N. gruberi***. **a**) Global methylation levels as quantified with EM-seq for each replicate of *A. castellanii* cells treated for 3 days with DMSO (control) and the cytidine analogues 5-Azacytidine, Decitabine and Zebularine. Black dots indicate the global methylation level, and red dots indicate the non-conversion rate for that library as estimated from the lambda spike-in genome. **b**) Distribution of methylation levels on hypermethylated regions fo the *A. castellanii* genome (merged genome bins with mCG ≥ 20%) for each treatment. **c**) Comparison of RNA-seq fold changes when comparing 5-Azacytidine treated *A. castellanii* cells against control and Decitabine treated samples against DMSO, as calculated by DEseq2. **d**) Number of differentially expressed genes (p adjusted value < 0.05) in the 5-

Azacytidine and the Decitabine treatments, classified as per methylation status and direction of change. **e**) Global methylation levels as quantified with EM-seq for each replicate of N gruberi cells treated for 3 days with DMSO (control) and the cytidine analogues 5-Azacytidine, Decitabine and Zebularine. Legend as in panel **a**. **f**) Comparison of RNA-seq fold changes when comparing 5-Azacytidine treated N. gruberi cells against control and Decitabine treated samples against DMSO, as calculated by DEseq2. **g**) Number of differentially expressed genes (p adjusted value < 0.05) in the 5-Azacytidine and the Decitabine treatments, classified as per methylation status and direction of change.

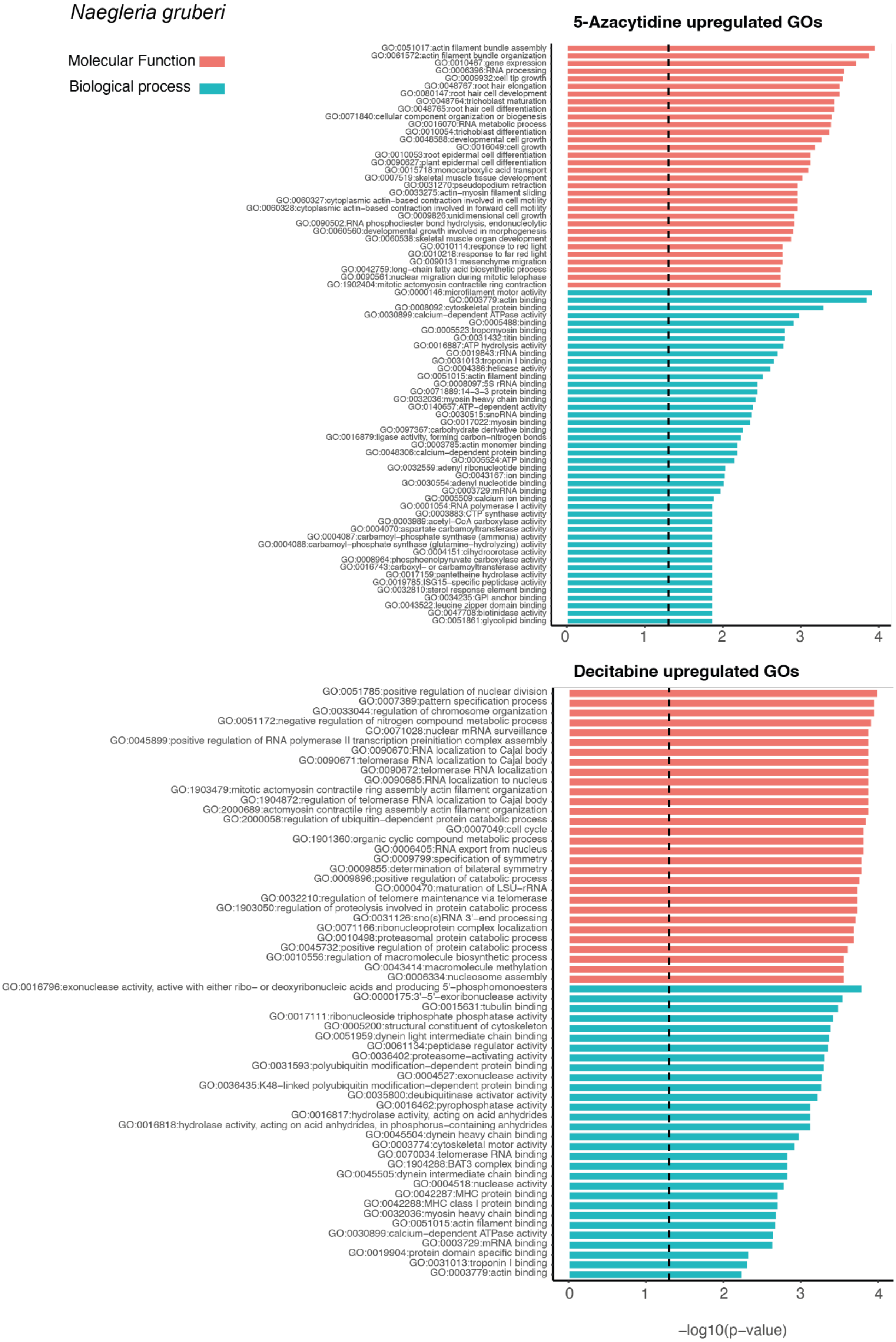


**Supplementary Figure 5. Gene Ontology enrichments of N. gruberi cytidine treated**

**response genes.** Gene ontology terms enriched for the differentially expressed genes upregulated upon 5-Azacytidine and Decitabine treatments. Enrichments computed with TopGO based on eggNOG-mapper annotations, shown for Molecular Function in red and Biological Process in green and sorted by p-value.

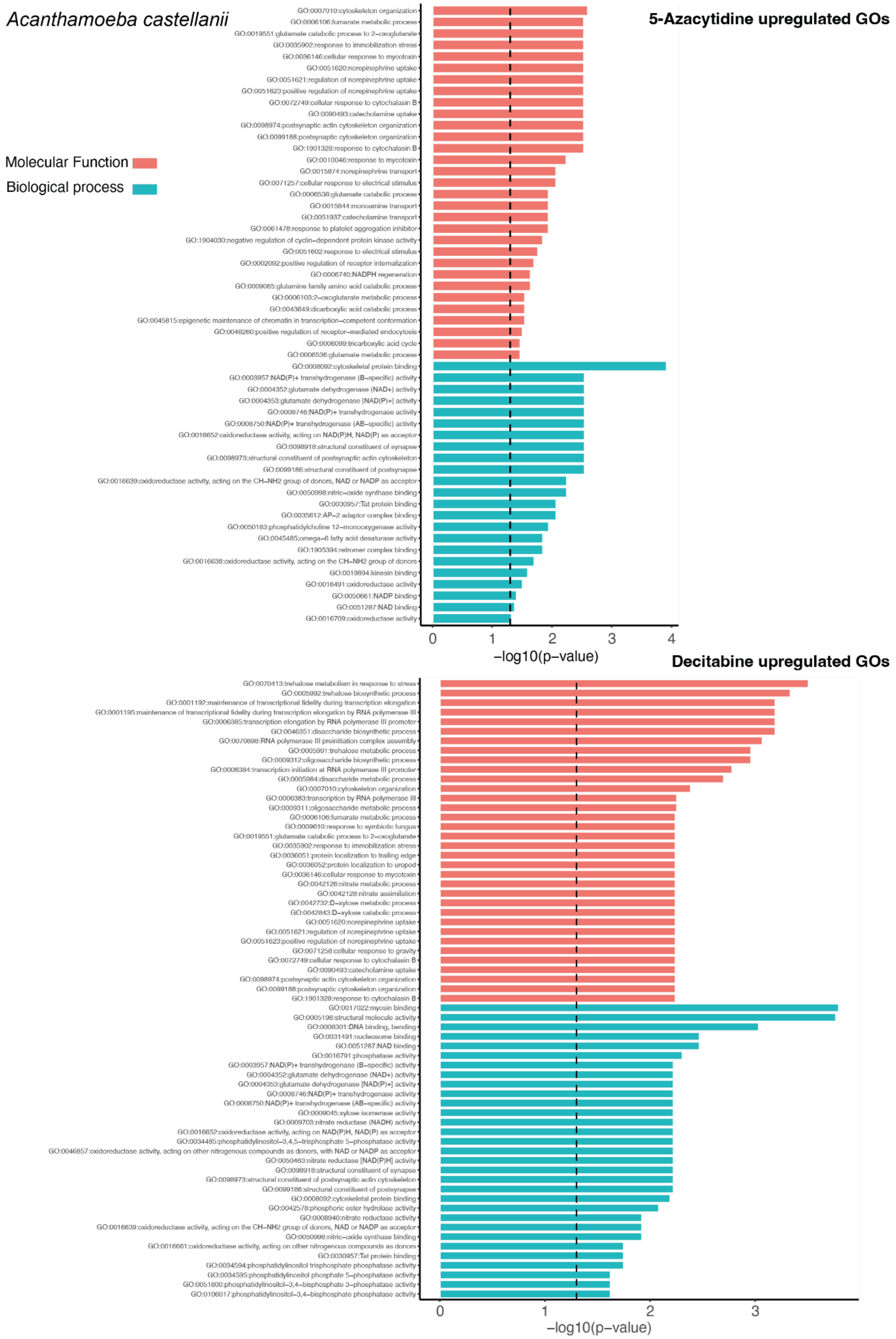
Acanthamoeba castellanii
5-Azacytidine upregulated GOs
Molecular Function
Biological process
GO:0007010:cytoskeleton organization
GO:0006106:fumarate metabolic process
GO:0019551:glutamate catabolic process to 2−oxoglutarate
GO:0035902:response to immobilization stress
GO:0036146:cellular response to mycotoxin
GO:0051620:norepinephrine uptake
GO:0051621:regulation of norepinephrine uptake
GO:0051623:positive regulation of norepinephrine uptake
GO:0072749:cellular response to cytochalasin B
GO:0090493:catecholamine uptake
GO:0098974:postsynaptic actin cytoskeleton organization
GO:0099188:postsynaptic cytoskeleton organization
GO:1901328:response to cytochalasin B
GO:0010046:response to mycotoxin
GO:0015874:norepinephrine transport
GO:0071257:cellular response to electrical stimulus
GO:0006538:glutamate catabolic process
GO:0015844:monoamine transport
GO:0051937:catecholamine transport
GO:0061478:response to platelet aggregation inhibitor
GO:1904030:negative regulation of cyclin−dependent protein kinase activity
GO:0051602:response to electrical stimulus
GO:0002092:positive regulation of receptor internalization
GO:0006740:NADPH regeneration
GO:0009065:glutamine family amino acid catabolic process
GO:0006103:2−oxoglutarate metabolic process
GO:0043649:dicarboxylic acid catabolic process
GO:0045815:epigenetic maintenance of chromatin in transcription−competent conformation
GO:0048260:positive regulation of receptor−mediated endocytosis
GO:0006099:tricarboxylic acid cycle
GO:0006536:glutamate metabolic process
GO:0008092:cytoskeletal protein binding
GO:0003957:NAD(P)+ transhydrogenase (B−specific) activity
GO:0004352:glutamate dehydrogenase (NAD+) activity
GO:0004353:glutamate dehydrogenase [NAD(P)+] activity
GO:0008746:NAD(P)+ transhydrogenase activity
GO:0008750:NAD(P)+ transhydrogenase (AB−specific) activity
GO:0016652:oxidoreductase activity, acting on NAD(P)H, NAD(P) as acceptor
GO:0098918:structural constituent of synapse
GO:0098973:structural constituent of postsynaptic actin cytoskeleton
GO:0099186:structural constituent of postsynapse
GO:0016639:oxidoreductase activity, acting on the CH−NH2 group of donors, NAD or NADP as acceptor
GO:0050998:nitric−oxide synthase binding
GO:0030957:Tat protein binding
GO:0035612:AP−2 adaptor complex binding
GO:0050183:phosphatidylcholine 12−monooxygenase activity
GO:0045485:omega−6 fatty acid desaturase activity
GO:1905394:retromer complex binding
GO:0016638:oxidoreductase activity, acting on the CH−NH2 group of donors
GO:0019894:kinesin binding
GO:0016491:oxidoreductase activity
GO:0050661:NADP binding
GO:0051287:NAD binding
GO:0016709:oxidoreductase activity
0
1
2
3
4
−log10(p−value)
Decitabine upregulated GOs
GO:0070413:trehalose metabolism in response to stress
GO:0005992:trehalose biosynthetic process
GO:0001192:maintenance of transcriptional fidelity during transcription elongation
GO:0001195:maintenance of transcriptional fidelity during transcription elongation by RNA polymerase III
GO:0006385:transcription elongation by RNA polymerase III promoter
GO:0046351:disaccharide biosynthetic process
GO:0070898:RNA polymerase III preinitiation complex assembly
GO:0005991:trehalose metabolic process
GO:0009312:oligosaccharide biosynthetic process
GO:0006384:transcription initiation at RNA polymerase III promoter
GO:0005984:disaccharide metabolic process
GO:0007010:cytoskeleton organization
GO:0006383:transcription by RNA polymerase III
GO:0009311:oligosaccharide metabolic process
GO:0006106:fumarate metabolic process
GO:0009610:response to symbiotic fungus
GO:0019551:glutamate catabolic process to 2−oxoglutarate
GO:0035902:response to immobilization stress
GO:0036051:protein localization to trailing edge
GO:0036052:protein localization to uropod
GO:0036146:cellular response to mycotoxin
GO:0042126:nitrate metabolic process
GO:0042128:nitrate assimilation
GO:0042732:D−xylose metabolic process
GO:0042843:D−xylose catabolic process
GO:0051620:norepinephrine uptake
GO:0051621:regulation of norepinephrine uptake
GO:0051623:positive regulation of norepinephrine uptake
GO:0071258:cellular response to gravity
GO:0072749:cellular response to cytochalasin B
GO:0090493:catecholamine uptake
GO:0098974:postsynaptic actin cytoskeleton organization
GO:0099188:postsynaptic cytoskeleton organization
GO:1901328:response to cytochalasin B
GO:0017022:myosin binding
GO:0005198:structural molecule activity
GO:0008301:DNA binding, bending
GO:0031491:nucleosome binding
GO:0051287:NAD binding
GO:0016791:phosphatase activity
GO:0003957:NAD(P)+ transhydrogenase (B−specific) activity
GO:0004352:glutamate dehydrogenase (NAD+) activity
GO:0004353:glutamate dehydrogenase [NAD(P)+] activity
GO:0008746:NAD(P)+ transhydrogenase activity
GO:0008750:NAD(P)+ transhydrogenase (AB−specific) activity
GO:0009045:xylose isomerase activity
GO:0009703:nitrate reductase (NADH) activity
GO:0016652:oxidoreductase activity, acting on NAD(P)H, NAD(P) as acceptor
GO:0034485:phosphatidylinositol−3,4,5−trisphosphate 5−phosphatase activity
GO:0046857:oxidoreductase activity, acting on other nitrogenous compounds as donors, with NAD or NADP as acceptor
GO:0050463:nitrate reductase [NAD(P)H] activity
GO:0098918:structural constituent of synapse
GO:0098973:structural constituent of postsynaptic actin cytoskeleton
GO:0099186:structural constituent of postsynapse
GO:0008092:cytoskeletal protein binding
GO:0042578:phosphoric ester hydrolase activity
GO:0008940:nitrate reductase activity
GO:0016639:oxidoreductase activity, acting on the CH−NH2 group of donors, NAD or NADP as acceptor
GO:0050998:nitric−oxide synthase binding
GO:0016661:oxidoreductase activity, acting on other nitrogenous compounds as donors
GO:0030957:Tat protein binding
GO:0034594:phosphatidylinositol trisphosphate phosphatase activity
GO:0034595:phosphatidylinositol phosphate 5−phosphatase activity
GO:0051800:phosphatidylinositol−3,4−bisphosphate 3−phosphatase activity
GO:0106017:phosphatidylinositol−3,4−bisphosphate phosphatase activity
0
1
2
3
−log10(p−value)

**Supplementary Figure 6. Gene Ontology enrichments of *A. castellanii* cytidine treated response genes.** Gene ontology terms enriched for the differentially expressed genes upregulated upon 5-Azacytidine and Decitabine treatments. Enrichments computed with TopGO based on eggNOG-mapper annotations, shown for Molecular Function in red and Biological Process in green and sorted by p-value.

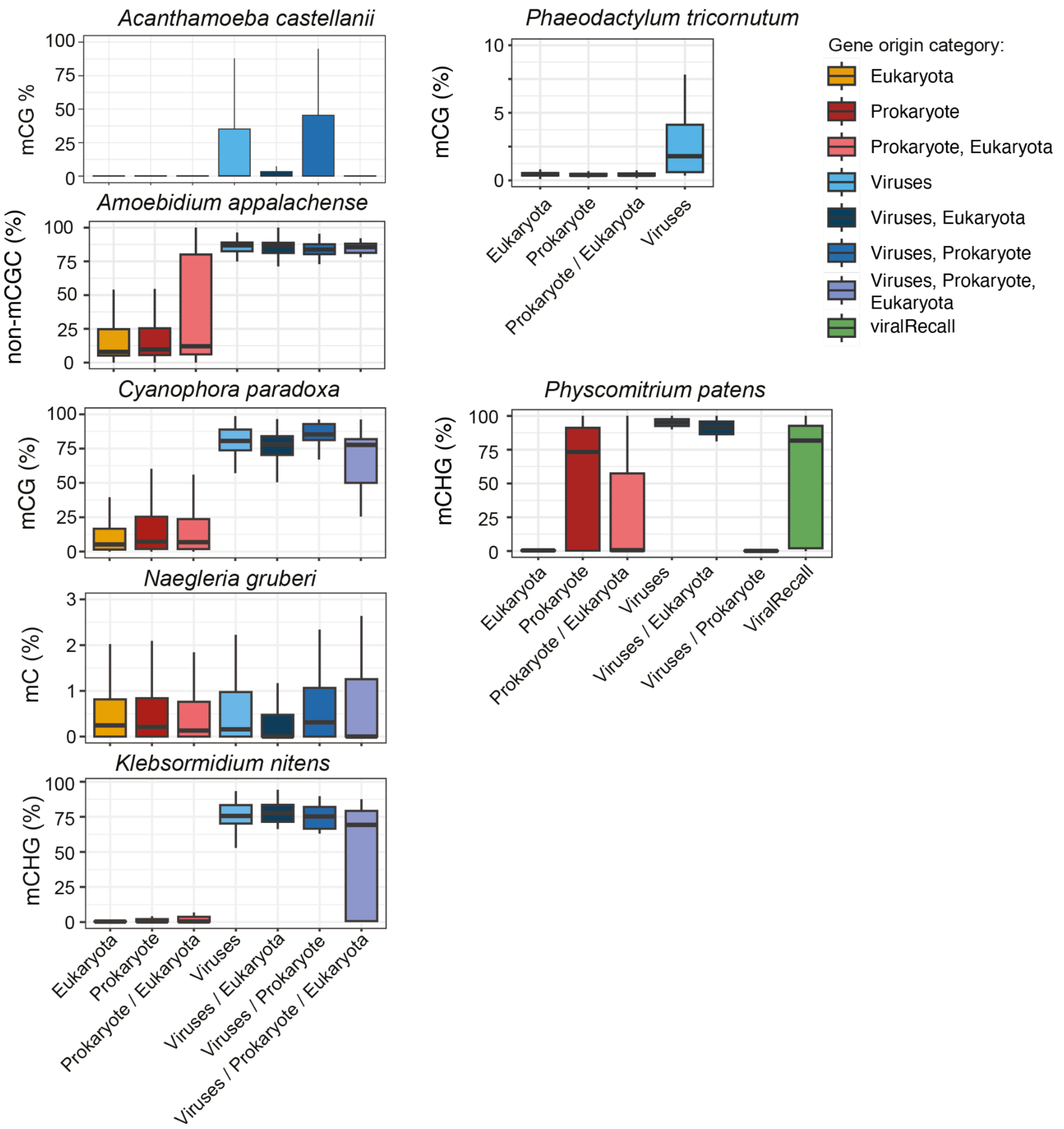


**Supplementary Figure 7. Methylation levels on genes by taxonomic origin.** Distribution of methylation levels for all genes in each given genome classified as per their best 10 hits against NCBI non redundant database. For *P. patens*, we ran ViralRecall as all viral genes have been masked from the latest Phypa V5 genome version. For *A. castellanii* methylated genes are defined as mCG ≥ 20%, in *Amoebidium appalachense / parasiticum* as non-mCGC methylation ≥ 70%, for *N. gruberi* mC ≥ 5%, for *C. paradoxa* mCG ≥ 50 %, for *K. nitens* and *P. patens* mCHG ≥ 10 %, for *P. tricornutum* mCG ≥ 1 %. White asterisks represent statistical enrichment (two-sided Fisher exact test < 0.01).
